# Shared texture-like representations underlie deep neural network alignment with human visual processing

**DOI:** 10.1101/2025.08.29.673066

**Authors:** Jessica Loke, Lynn K.A. Sörensen, Iris I.A. Groen, Natalie Cappaert, H. Steven Scholte

## Abstract

Deep neural networks (DNNs) excel at predicting neural responses across the visual hierarchy ^1–5^, a success widely interpreted as evidence of shared object recognition computations ^6,7^. Yet improving DNN object recognition accuracy does not reliably increase neural predictivity ^8,9^, and even untrained networks predict brain responses above chance ^9–11^. This disconnect suggests that object recognition may not drive DNN-brain alignment. Texture-like statistics are represented in both DNNs and mid-level visual cortex ^12–17^. In natural images, these statistics are carried by objects and backgrounds, shaping representations and recognition in both systems ^18–21^. Does DNN-brain alignment reflect shared sensitivity to object-related information or texture-like statistics? To dissociate these factors, we recorded EEG from 57 participants viewing natural scenes, texture-synthesized images preserving local statistics while disrupting global form and object-only images with backgrounds removed. If alignment reflects texture-like statistics, it should peak for texture-synthesized images. If it reflects object-related processing, alignment should be strongest for natural and object-only conditions, which preserve object information. We compared EEG responses with DNN activations via weighted representational similarity analysis ^22,23^. Texture-synthesized images yielded the strongest DNN-EEG alignment, peaking in early responses (<200 ms) and explaining up to ∼85% of noise-ceiling-normalized explainable variance versus ∼44% for natural and ∼55% for isolated objects. Crucially, object categories were more decodable for natural and object-only images than texture-synthesized images, yet these object-rich conditions showed weaker alignment. This dissociation reveals that DNNs capture the texture-statistical component of early visual responses while failing to explain later, object-related variance.

**Highlights:**

- Texture-synthesized images yield the strongest DNN-brain alignment across DNNs.
- Object categories decode better from object-rich than texture-synthesized EEG.
- DNN-brain alignment reflects shared texture statistics, not object-related processing.

## Results and discussion

DNNs predict neural responses across the visual hierarchy ^1–5^. This is widely interpreted as evidence of shared object recognition computation ^6,7^. Yet improving DNN object recognition accuracy does not reliably increase neural predictivity ^8,9^, and even untrained networks predict brain responses above chance ^9–11^. Here, we test whether alignment instead reflects shared sensitivity to local, texture-like statistics (which we define throughout as global summaries of local image statistics); these are the same features underlying DNNs’ texture bias ^24–27^ which are also well-represented in early and mid-level visual cortex ^12–17^. That early and mid-level cortex encodes these statistics is established; our question is not whether the brain represents them but which component of neural responses the alignment between DNNs and brains reflects. If DNN-brain alignment arises from this shared sensitivity rather than from shared object recognition computation, alignment should be highest for texture-synthesized images, which isolate these statistics from global form.

Importantly, real-world objects are often embedded within complex backgrounds; these backgrounds have been shown to shape neural responses, recognition accuracy, and influence the degree to which both humans and DNNs prioritize background over object information ^18–21,28^. Because objects and backgrounds inherently co-occur in the real world, isolating their respective contributions requires decoupling them.

Yet no study has directly contrasted texture-synthesized, natural, and object-only images to determine which visual component drives DNN-brain alignment. To close this gap, we recorded EEG from 57 participants viewing three image conditions in an RSVP paradigm (100 ms/image): original natural scenes, texture-synthesized counterparts generated by VGG-19 conv1_1 Gram-matrix synthesis ^29^ (local statistics preserved, global form disrupted), and object-only images (background texture removed, object form retained; Figure 1A–B). We compared DNN activations from five architectures (AlexNet, VGG-16, ResNet-18, ResNet-50, ViT-B/16) to time-resolved EEG representational dissimilarity matrices (RDMs) using weighted representational similarity analysis (RSA) ^22,30^. The overall alignment was summarized as the noise-ceiling-normalized area under the curve (AUC; STAR Methods; Figure 2A–B). Because we recorded EEG to stimuli that selectively preserve texture-like image statistics or object form, and can derive DNN features for each, we can ask not only whether DNN features align with neural responses but which image properties that alignment reflects. We addressed this by testing cross-condition generalization of DNN features (Figure 3) and decoding object category information from EEG to dissociate representational alignment from object-related neural signals (Figure 4).

**Figure 1.**
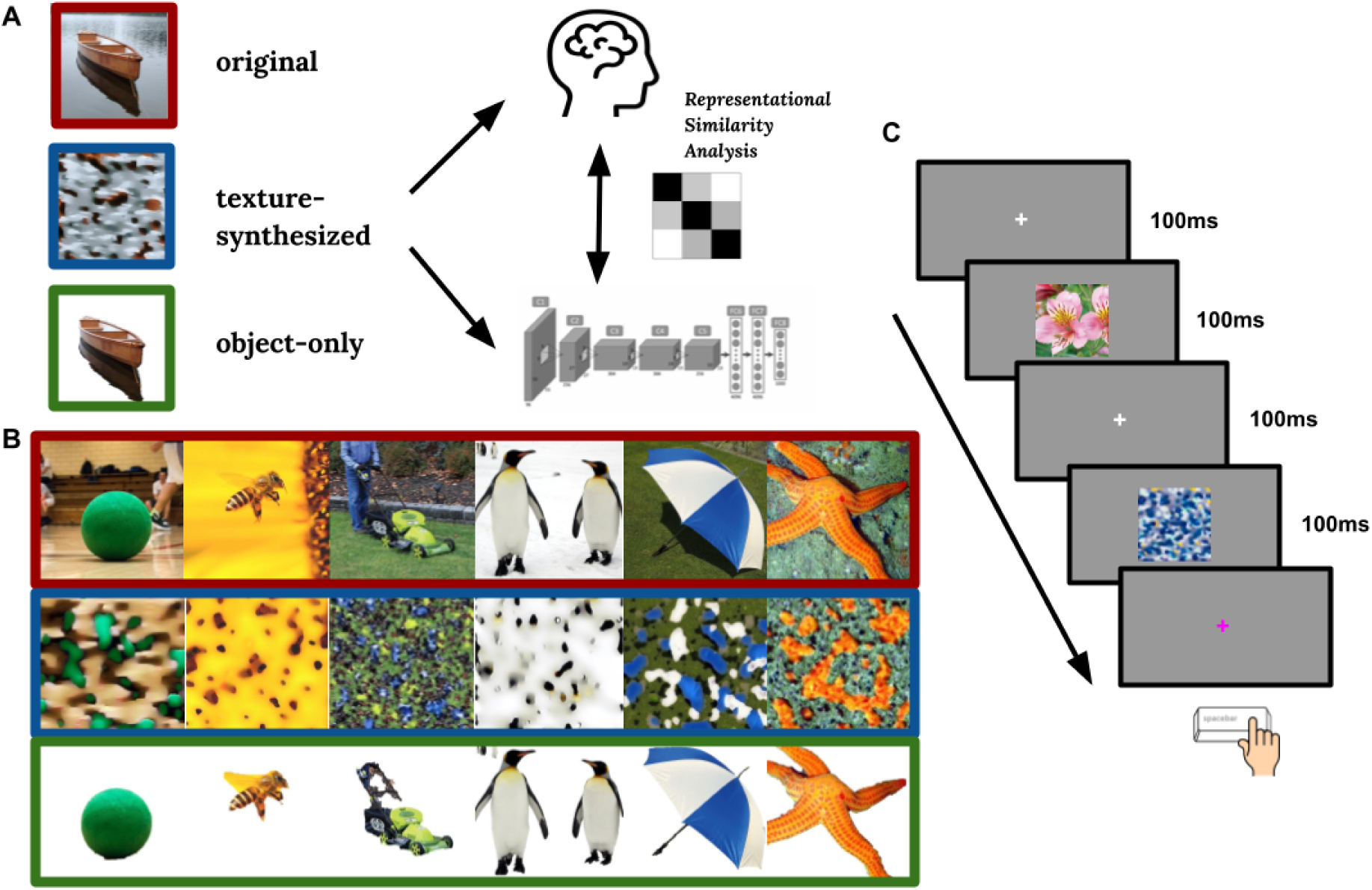
Experimental design and images used. (a) Original natural images (texture + object + background statistics), texture-synthesized images (texture statistics only; global form disrupted), and object-only images (texture + object form; background removed) were presented to human participants and deep neural networks (DNNs). Representational dissimilarity matrices (RDMs) were constructed from EEG responses and DNN activations (five architectures, all layers) and compared using weighted representational similarity analysis (RSA combined with ridge regression), cross-validated over held-out participants and stimuli. (b) Examples of original THINGS images (top row), texture-synthesized counterparts (middle row), object-only versions (bottom row). (c) Participants viewed stimuli in a rapid serial visual presentation paradigm where images were presented at 100 ms with a 200 ms stimulus-onset asynchrony while performing an orthogonal fixation-color detection task.

**Figure 2.**
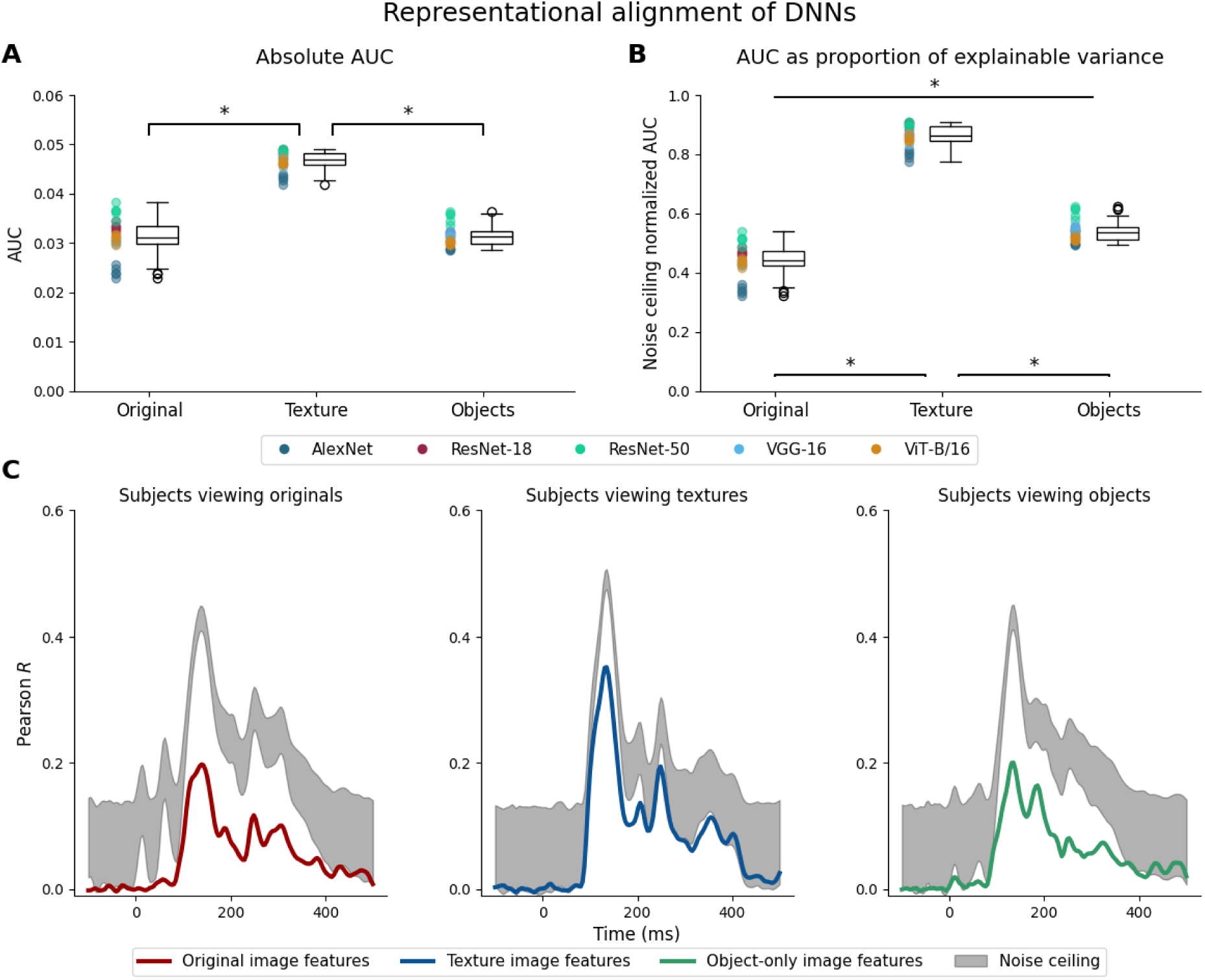
Representational alignment of DNNs on matching stimuli. (A & B) Representational alignment is quantified as area under the curve (AUC) of Pearson *r* across 0-500 ms post-stimulus. Each colored dot is a model initialization, with colors grouping DNN architectures. Box plots show the distribution across all architectures and initializations. Subplot A shows the representational alignment in absolute AUC values, whereas B shows the AUC values normalized by the upper bounds of the noise ceiling. In both A & B subplots, pairwise Wilcoxon signed-rank tests show that DNNs’ representational alignment for texture-synthesized images is significantly higher than alignment for original and object-only images, while no significant differences were observed between the original and object-only conditions using absolute AUC values. This enhanced representational alignment for texture-synthesized images across all DNN architectures suggests that texture-like image statistics drive the representational alignment between DNNs and human visual processing. Asterisks indicate statistically significant differences (*p* <. 01, Bonferroni-corrected). (C) The full time course of the representational alignment averaged across all DNN architectures. Gray shaded regions represent noise ceiling bounds.

**Figure 3.**
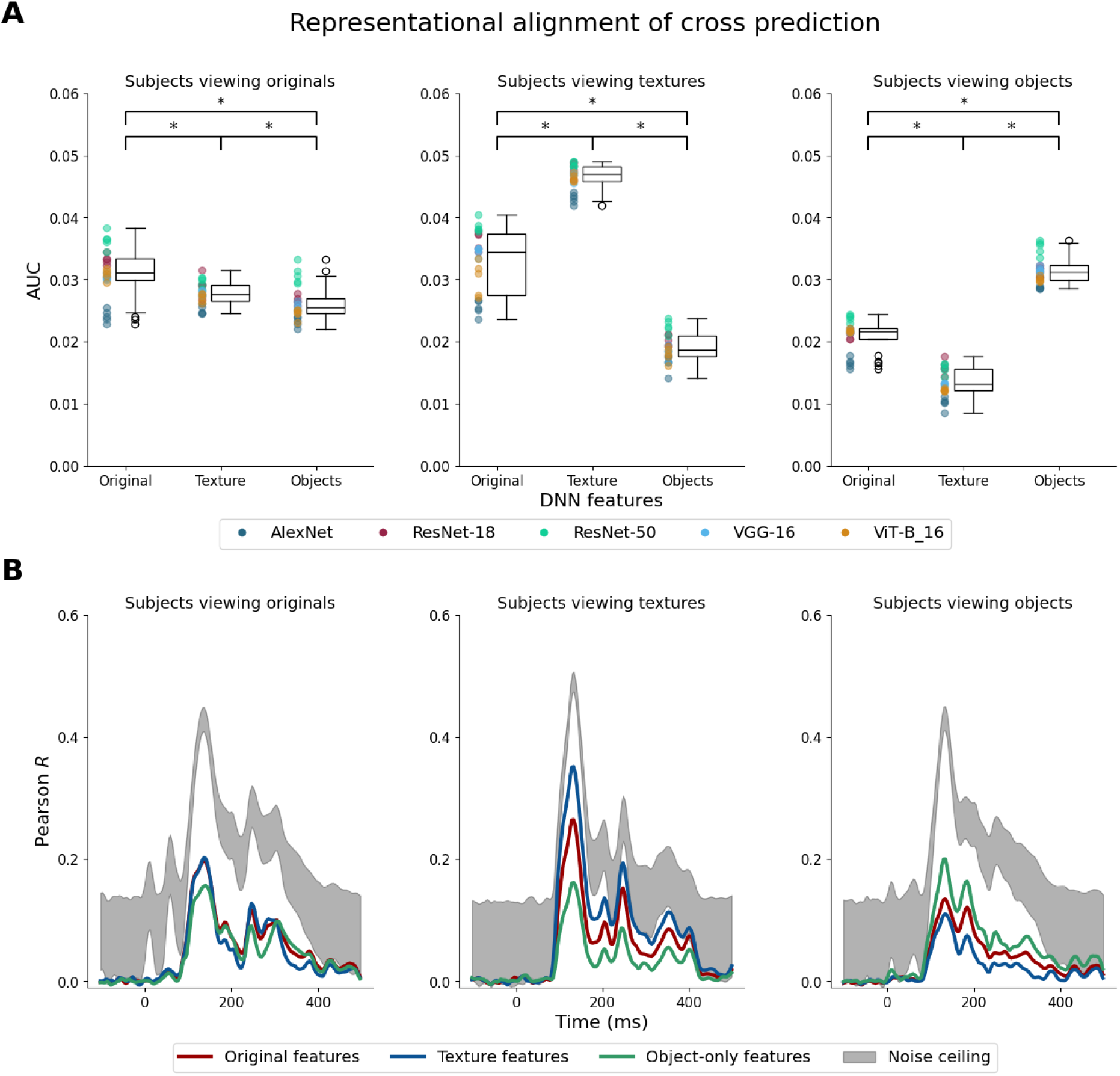
Cross-condition representational alignment reveals DNNs selectively capture variance in neural responses related to local image statistics. (A) Representational alignment (AUC) when DNN features from one stimulus condition are used to predict EEG responses to another. Each panel shows EEG responses to one stimulus condition (left: original; middle: texture-synthesized; right: object-only). Each point represents a model initialization with colors corresponding to different architectures. Box plots show the distribution of AUC values across architectures. For EEG responses to original images (left panel), original features significantly outperformed texture-synthesized features in predicting neural responses. However, texture-synthesized features achieved nearly comparable performance, demonstrating that DNNs predominantly model the texture-statistical components of neural responses. Asterisks indicate statistically significant differences (*p* <. 01, Bonferroni-corrected). (B) The full time course of the cross-condition alignment, averaged across all DNN architectures. The cross-prediction is performed using features from different image conditions (colored lines: red - original features, blue - texture features, green - object features). Gray shaded regions represent noise ceiling bounds.

**Figure 4.**
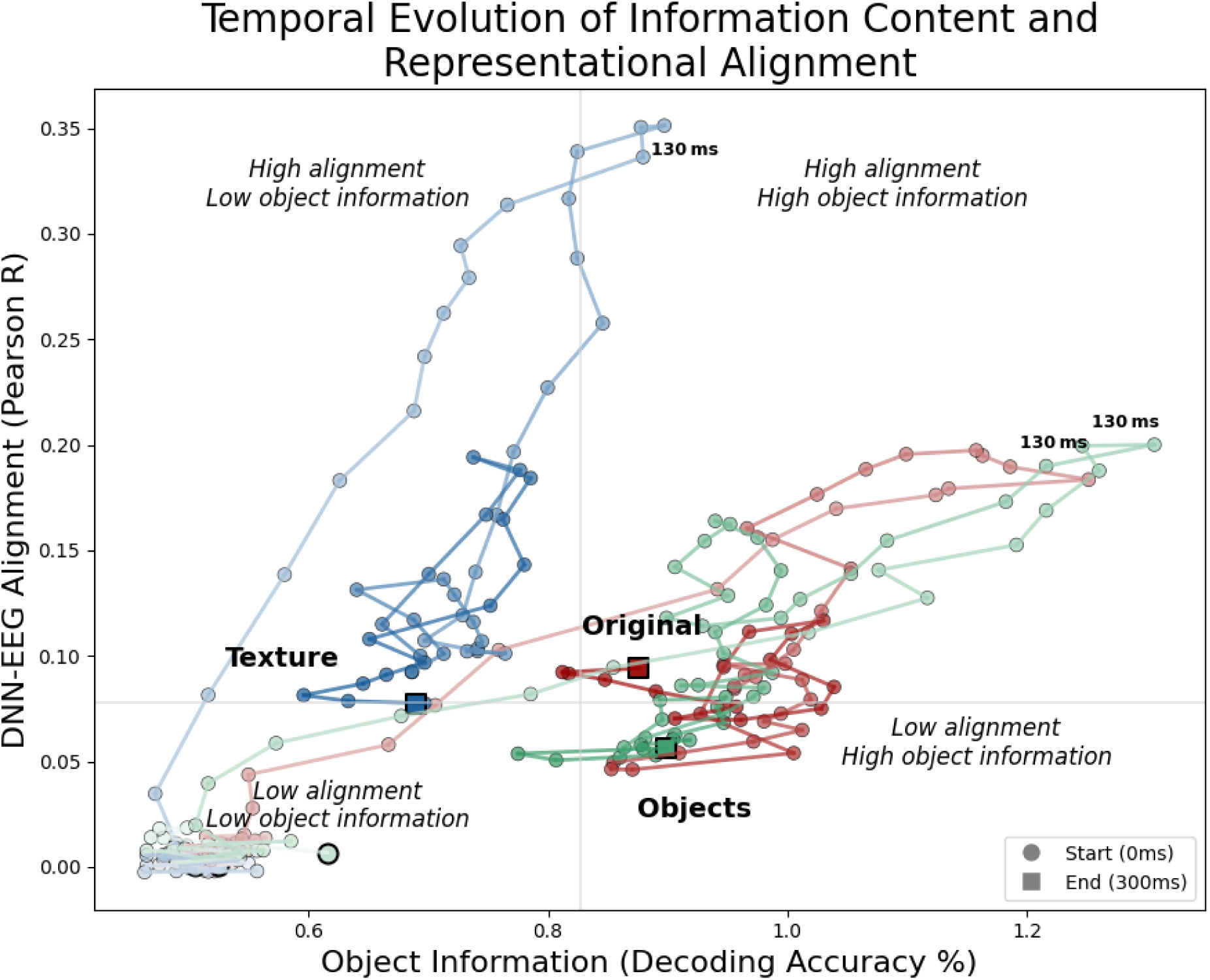
Temporal evolution of object information and representational alignment. Each trajectory plots object information (x-axis, measured by decoding accuracy) against DNN-EEG alignment (y-axis, Pearson *r*) from stimulus onset to 300ms. Colors represent different image conditions: original (red), texture-synthesized (blue), object-only (green). Lighter colors mark earlier latencies, darker colors mark later latencies. Circles mark trajectory starts (0 ms) and squares mark ends (300 ms). The trajectories reveal a clear dissociation between object information and DNN-EEG alignment – alignment reaches its maximum when object information is minimal (texture-synthesized condition), whereas conditions rich in object form yield higher object information without a corresponding boost in alignment. This pattern demonstrates that DNN-brain alignment primarily tracks neural variance related to global summaries of local image statistics, while leaving object-related components comparatively unexplained.

### Texture-synthesized images maximize DNN-EEG representational alignment

Texture-synthesized images produced the highest DNN-EEG alignment (Friedman test: χ²(2)=37.68, p<.001; Figure 2A). This advantage was consistent across all five architectures despite their range of ImageNet recognition accuracy (AlexNet ∼51% to ViT-B/16 ∼81%; Figure S1), indicating that recognition performance does not drive alignment. Pairwise Wilcoxon signed-rank tests confirmed that alignment for texture-synthesized images exceeded both original (W=0, r=0.87, p<.001) and object-only alignment (W=0, r=0.87, p<.001; Bonferroni-corrected). This advantage was not an artifact of differences in the overall distribution of dissimilarity values as randomly scrambling the model RDMs abolished the alignment across all conditions (Figure S2). After normalization by the noise ceiling, texture-synthesized features explained up to ∼85% of maximum explainable variance versus ∼44% for natural scenes and ∼55% for object-only images (Figure 2B). Alignment peaked within 100–200 ms of stimulus onset across all conditions, consistent with the feedforward sweep through early visual areas. Notably, the noise ceiling was higher after 200 ms for natural and object-only conditions than for texture-synthesized images (Figure 2C), indicating that cross-participant consistency for object-related signals emerged in this later window. These later signals were less well explained by the DNNs as compared to earlier ones, suggesting a failure to account for object-related neural variance.

### Texture-synthesized features generalize to natural scenes better than object-only features

If shared sensitivity to local image statistics drives DNN-EEG alignment, features extracted from texture-synthesized images should generalize to natural-image EEG as well as or better than object-only features. Texture-synthesized features preserve full image local statistics, whereas object-only images retain object-related information but remove background and thus much of the contextual statistical structure. We used a cross-condition analysis (Figure 3) to understand the relative importance of texture versus object-related information for DNN-EEG alignment. When predicting EEG responses to natural scenes (Friedman χ²(2)=30.48, p<.001), texture-synthesized features achieved an identical peak alignment to original features (Pearson r=0.20 at 138 ms), although original features showed significantly higher AUC across 0-500ms (W=17, p<.001, r=0.78). Original features also outperformed object-only features (W=1, p<.001, r=0.87). Strikingly, texture-synthesized features outperformed object-only features even for natural image EEG (W=42, p=.002, r=0.65), a counterintuitive result given that natural and object-only images are perceptually more similar to each other than either is to texture-synthesized images.

In the remaining two conditions, alignment was instead dominated by the matching features (texture features for texture-synthesized EEG, object-only features for object-only EEG); critically, even this best-matched alignment was substantially lower for object-only than for texture-synthesized images (Figure 3).

This indicates that removing background texture reduces overall DNN-EEG correspondence rather than isolating a stronger object-related signal.

Together, these results show that DNN-EEG alignment is carried by texture-like image statistics instead of object-related information. Texture features generalize to natural scenes, which share those statistics, but not to isolated objects, where object form is preserved while much of the background/contextual statistical structure is removed.

### Object information is decodable from EEG but does not drive alignment with DNN

To verify that EEG captured object-related information, we decoded the object category from single-trial EEG patterns at each time sample (STAR Methods). Despite a brief 100 ms presentation, all conditions yielded above-chance decoding across 200 object categories, with the full time course of object decoding shown in Figure S4. If DNN-brain alignment reflects shared object-related representations, conditions demonstrating more object-related information should also show stronger alignment. Intriguingly, the conditions with weaker DNN-EEG alignment (natural and object-only condition) contained the most object information (highest decoding). To investigate this apparent dissociation between object information and DNN-EEG alignment, we traced the co-evolution of object information (decoding accuracy) and DNN-EEG alignment (Pearson’s *r*) from stimulus onset to 300 ms (Figure 4). A shared object recognition account would predict that natural and object-only trajectories show the highest DNN-EEG alignment as they have richer object information. Instead, natural and object-only conditions move into the high-object-information/low-alignment quadrant, while texture-synthesized moves sharply into the high-alignment/low-object-information quadrant. This directly demonstrates that DNN-brain alignment arises through texture-like image statistics, not object-related processing. This dissociation also explains why boosting object recognition accuracy does not consistently improve neural predictivity ^8,9^. Improvements to representations DNNs don’t share with the brain leave the alignment unchanged.

### DNN-brain alignment is dominated by early feedforward processing

Time-resolved alignment (Figure 3B) peaks within 100–200 ms of stimulus onset, consistent with afferent feedforward activity in V1–V4 ^31,32^. After 200 ms, the window dominated by recurrent processing ^31^, the noise ceiling is higher in the natural and object-only viewing conditions than in the texture-synthesized viewing condition. Although DNN-EEG alignment in all three conditions is lower after the early peak between 100 and 200 ms, alignment in the texture-synthesized condition approaches the lower noise ceiling bound, whereas a larger gap persists for the original and object-only viewing conditions. This pattern indicates that later object-related neural signals in the natural and object-only conditions are less well explained by current feedforward architectures. Overall, these results are consistent with DNN-EEG alignment being dominated by early feedforward encoding.

### Shared texture sensitivity, not object processing, drives DNN-brain alignment

Earlier work established that DNNs are more texture-biased than humans behaviorally ^24^ and that early visual cortex encodes texture-like statistics ^12–17^; whether this shared sensitivity drives DNN-brain representational alignment had not been tested.

DNN-brain alignment is usually read as evidence of shared object recognition; our EEG results indicate it instead reflects a shared sensitivity to early, texture-like image statistics rather than object recognition computations.

This alignment is concentrated in the initial feedforward processes, before further processing transforms these local statistics into stable object percepts ^31,33^. Because neural predictivity declines as object-specific neural signals peak, high DNN–brain alignment is better understood as a signature of early statistical extraction than of the more complex processing required for human object recognition.

These findings might appear at odds with the human behavioral shape bias ^24^: if the visual system ultimately prioritizes shape for object recognition, strong DNN-EEG alignment driven by texture statistics may seem paradoxical. Yet the shape bias is a behavioral readout of the full processing hierarchy ^13,17^. Current DNNs primarily excel at capturing this early, feedforward stage of visual processing rather than the full processing chain. Texture-dominated alignment at this early stage is still consistent with a system that ultimately behaves with a shape bias.

Our texture-synthesized stimuli preserve only local image statistics, with global object form disrupted. That they nonetheless produce the strongest DNN-EEG alignment helps explain why even untrained networks achieve above-chance neural prediction scores ^9–11^. If alignment reflects shared sensitivity to texture-like statistics rather than learned object recognition, neither architectural sophistication nor training is required to capture it. Advancing models of object-centered vision will therefore require architectures that capture the later, object-related processing current DNNs miss, for instance by explicitly representing global shape.

### Limitations and conclusion

Several limitations bound these conclusions. First, the 100 ms RSVP paradigm may bias processing toward feedforward representations as participants were not required to recognize the objects. Task demands can modulate the magnitude and temporal profile of early visual responses without necessarily determining their overall representational structure. Nevertheless, our orthogonal fixation task may have favored a more feedforward processing regime. We therefore cannot assume that the texture advantage would generalize unchanged to explicit object recognition tasks. Second, our texture synthesis simultaneously scrambles background and object textures, preventing dissociation of their contributions. Future studies could vary texture statistics across identical objects to clarify the relative contributions of object identity and texture features to neural representational similarity. Third, our 17 posterior electrodes also favor superficial visual sources over deeper regions such as anterior inferotemporal cortex; including all electrodes preserved the ranking (Figure S5), but EEG’s limited sensitivity to deep sources leaves our object-related measure partial.

Finally, by eliminating global form, texture-synthesized stimuli also remove semantic content, likely attenuating top-down feedback after 200 ms, such that the high alignment for these stimuli may partly reflect reduced top-down modulation in addition to shared feedforward texture coding. However, because the texture advantage in DNN-EEG alignment peaks within 100-200ms, before recurrent processing dominates, reduced top-down modulation does not account for the effect. Future work could quantify this top-down contribution using stimuli that preserve semantic content.

Together, these results show that the correspondence between current DNNs and early human visual responses is carried by texture-like image statistics, while the object-related signals present in those responses go uncaptured by the networks. Advancing models of object-centered vision will therefore require architectures that capture this later, object-related processing.

## Resource Availability

### Lead contact

Further information and requests for resources should be directed to and will be fulfilled by the lead contact, H. Steven Scholte.

### Materials availability

This study did not generate new reagents or materials.

### Data and code availability

Raw and processed EEG data and all analysis code will be deposited in a public repository and made available upon publication.

### Author Contributions

Conceptualization: J.L. and H.S.S. Methodology: J.L. Software: J.L. Formal analysis: J.L. and L.K.A.S. Investigation: J.L. Resources: H.S.S. and N.C. Data curation: J.L. Writing – original draft: J.L. Writing – review and editing: J.L., L.K.A.S., I.I.A.G., N.C., and H.S.S. Visualization: J.L. Supervision: H.S.S. Funding acquisition: H.S.S. and N.C.

### Declaration of Interests

The authors declare no competing interests.

## STAR★Methods

### Key Resources Table

| REAGENT or RESOURCE | SOURCE | IDENTIFIER |
| --- | --- | --- |
| Deposited data |  |  |
| Raw and analyzed EEG data | This paper | [repository upon publication] |
| Datasets |  |  |
| THINGS image dataset | Hebart et al., 2019 <sup>34</sup> | https://things-initiative.org |
| ImageNet ILSVRC 2012 | Russakovsky et al., 2015 <sup>35</sup> | https://image-net.org |
| Software and algorithms |  |  |
| Custom analysis code | This paper | [GitHub upon publication] |
| Python 3.12 | Python Software Foundation | https://www.python.org; RRID: SCR_008394 |
| MNE-Python | Gramfort et al., 2013 <sup>36</sup> | https://mne.tools; RRID: SCR_005972 |
| PyTorch v2.0.0 | Meta AI | https://pytorch.org; RRID: SCR_018536 |
| torchvision | PyTorch contributors | https://pytorch.org/vision |
| scikit-learn | Pedregosa et al., 2011 <sup>37</sup> | https://scikit-learn.org; RRID: SCR_002577 |
| SciPy v1.15.2 | Virtanen et al., 2020 <sup>38</sup> | https://scipy.org; RRID: SCR_008058 |
| NumPy | Harris et al., 2020 <sup>39</sup> | https://numpy.org; RRID: SCR_008633 |
| Background Remover v0.2.7 | Nader et al., 2023 <sup>40</sup> | https://github.com/nadermx/backgroundremover |
| Other |  |  |
| BioSemi ActiveTwo (64-channel) | BioSemi | https://www.biosemi.com |
| VGG-19 pretrained weights<br>(texture synthesis) | Simonyan and<br>Zisserman,<br>2014 <sup>41</sup> | https://pytorch.org/vision/stable/<br>models.html |
| ViT-B/16 pretrained weights<br>(ViT_B_16_Weights.DEFAULT) | PyTorch<br>contributors |  |

### Experimental Model and Subject Details

Fifty-seven participants (45 female, 12 male; ages 18–28) were recruited from the University of Amsterdam. Five were excluded (incorrect triggers n=1, premature termination n=1, noisy EEG n=3), leaving 52 in the final analysis. The study was approved by the ethical review board of the Faculty of Social and Behavioral Sciences, University of Amsterdam (FMG-3925_2023). All participants provided written informed consent and received course credit.

### Method Details

#### Stimuli

The 200 base images used were the held-out test set of THINGS-EEG2 ^10^, which contains one image from each of 200 object concepts. The remaining 1,654 concepts formed the THINGS-EEG2 training partition. The THINGS-EEG2 training and test split was pseudo-random and preserved the proportions of the 27 higher-level THINGS categories. We adopted the same test partition to align our stimuli with that benchmark. Texture-synthesized versions were generated from white noise by matching VGG-19 conv1_1 Gram matrices ^29^, minimizing the mean-squared Gram-matrix distance to preserve summaries of local statistics while disrupting global form. Object-only images were produced using automatic background removal Background Remover v0.2.7, followed by manual correction when necessary, with backgrounds replaced by white. Unlike texforms ^17^, which preserve coarse object form through localized spatial pooling, conv1_1 Gram-matrix synthesis removes substantially more global contour and layout information.

#### Experimental design and RSVP paradigm

Participants viewed original natural scenes, texture-synthesized images, and object-only images in a RSVP paradigm (Figure 1C). Stimuli from all three image types were randomly intermixed within each sequence, and trial order was independently randomized for each participant. A fixation cross was presented for 100 ms, followed by an image for 100ms. Successive image onsets were separated by 200ms. On 3% of fixation periods, the cross turned from white to pink; participants pressed the spacebar whenever this occurred, no image was presented afterward. Each of the 600 unique stimuli was presented 20 times (12,000 trials in total). A block consisted of 120 trials followed by a 15-second break. After five blocks, participants received a 90-second break. All participants maintained >90% accuracy on the fixation-color detection task.

#### EEG acquisition and preprocessing

EEG was recorded at 1024 Hz using 64-channel BioSemi ActiveTwo with a 10–10 cap layout. The data were re-referenced offline to the average of the left and right mastoid electrodes ^42^. We chose mastoid referencing because the main analyses focused on posterior visual electrodes, for which average referencing can alter the estimated scalp distribution when electrode coverage is spatially uneven ^42^. ICA was fitted on 1–30 Hz filtered data to identify artifacts through visual inspection – two ocular artifact components per participant were removed. ICA weights were applied to 0.1–30 Hz filtered data. Data was segmented into epochs from −100 to 500 ms relative to stimulus onset and baseline-corrected using the −100 to 0 ms pre-stimulus interval. No epochs were rejected. Because images were presented with a 200 ms stimulus onset asynchrony, this pre-stimulus interval may contain residual activity from the preceding stimulus. However, image order was randomized independently for each participant and image types were fully intermixed, so residual activity from the preceding image was not systematically related to the identity or condition of the current image. Epochs were downsampled to 256 Hz.

#### Deep neural network models

AlexNet, VGG-16, ResNet-18, and ResNet-50 were each trained from scratch on ImageNet ILSVRC 2012 with five random seeds (PyTorch v2.0.0). AlexNet was trained with batch size 512, learning rate 0.01, AdamW optimizer, and weight decay 1e-4 for 80 epochs. VGG-16 and ResNet-50 were trained with batch size 768, learning rate 0.005, AdamW optimizer, and weight decay 1e-4 for 80 epochs. ResNet-18 was trained with batch size 1024, learning rate 0.005, AdamW optimizer, and weight decay 1e-4 for 80 epochs. Batch size and learning rate were allowed to vary across architectures as these hyperparameters are architecture-dependent and were selected to support stable training within GPU-memory constraints. ViT-B/16 used one initialization of pretrained torchvision weights (ViT_B_16_Weights.DEFAULT). ViT-B/16 was not trained from scratch: vision transformers lack the inductive biases of convolutional networks and require large-scale pretraining or specialized augmentation and regularization schedules to train well on ImageNet, so we used the established torchvision weights rather than the simpler from-scratch regime applied to the convolutional models. Activations were extracted from all convolutional, pooling, and fully connected layers per condition. For ViT-B/16, activations were extracted from the convolutional patch-projection layer (`conv_proj; 14 x 14` patch embeddings) and from the multi-head self-attention and MLP/feedforward outputs of all 12 transformer encoder blocks.

#### Representational similarity analysis

At each time point, the response to each image was represented as a vector of amplitudes across 17 posterior electrodes (Oz, O1, O2, POz, PO3, PO4, PO7, PO8, Pz, P1–P8), selected as they primarily capture activity from visual processing areas, and averaged across the 20 repetitions of that image. Pairwise cosine distances between these vectors across all 200 images yielded a 200 × 200 EEG RDM per participant, condition, and time point (electrodes were not averaged; their amplitudes defined the response vector); only the upper triangle was used ^43^.

DNN RDMs were computed using the same cosine distance metric, applied to layer activations for each condition separately. Weighted RSA used Ridge regression (`sklearn RidgeCV, α` ∈ [10^−1^, 10^5^]) to regress EEG RDMs onto all DNN layer RDMs simultaneously per architecture. Cross-validation was stratified over 15 held-out participants and 100 held-out stimuli (50 repeated folds). Performance was quantified as Pearson *r* between predicted and observed held-out RDMs, summarised as AUC (trapezoidal integration, 0–500 ms).

#### Noise ceiling estimation

Noise ceilings were computed separately per image condition following Storrs et al. ^30^ and Nili et al. ^22^. Upper bound: correlation of each test participant’s RDM with the mean across all participants. Lower bound: correlation with the mean across training participants only. AUC was normalized by the upper noise ceiling AUC.

#### Object decoding analysis

A multi-class SVM was trained per participant, condition, and time sample to classify 200 object categories (chance=0.5%) using single-trial EEG patterns from 17 posterior electrodes. For each image, the 20 repetitions were randomly split 70/30 into a training set (14 repetitions) and a test set (6 repetitions); trials were not averaged across repetitions. Single-trial patterns were z-scored (`StandardScaler`) before classification (`RBF kernel`, C=1.0, γ=1/17).

### Quantification and Statistical Analysis

Friedman tests detected main effects of image condition; Wilcoxon signed-rank post hoc tests with Bonferroni correction assessed pairwise differences (α=.01). All tests were performed in Python 3.12 with SciPy v1.15.2. Sample size: 52 participants; 21 model initializations across five architectures.

## Notes

### Competing Interest Statement

The authors have declared no competing interest.

### Summary of Updates

This version of the manuscript has been revised from an empirical paper format to a shorter report format.

