## Supplemental Figures for "Shared texture-like representations underlie deep neural network alignment with human visual processing"

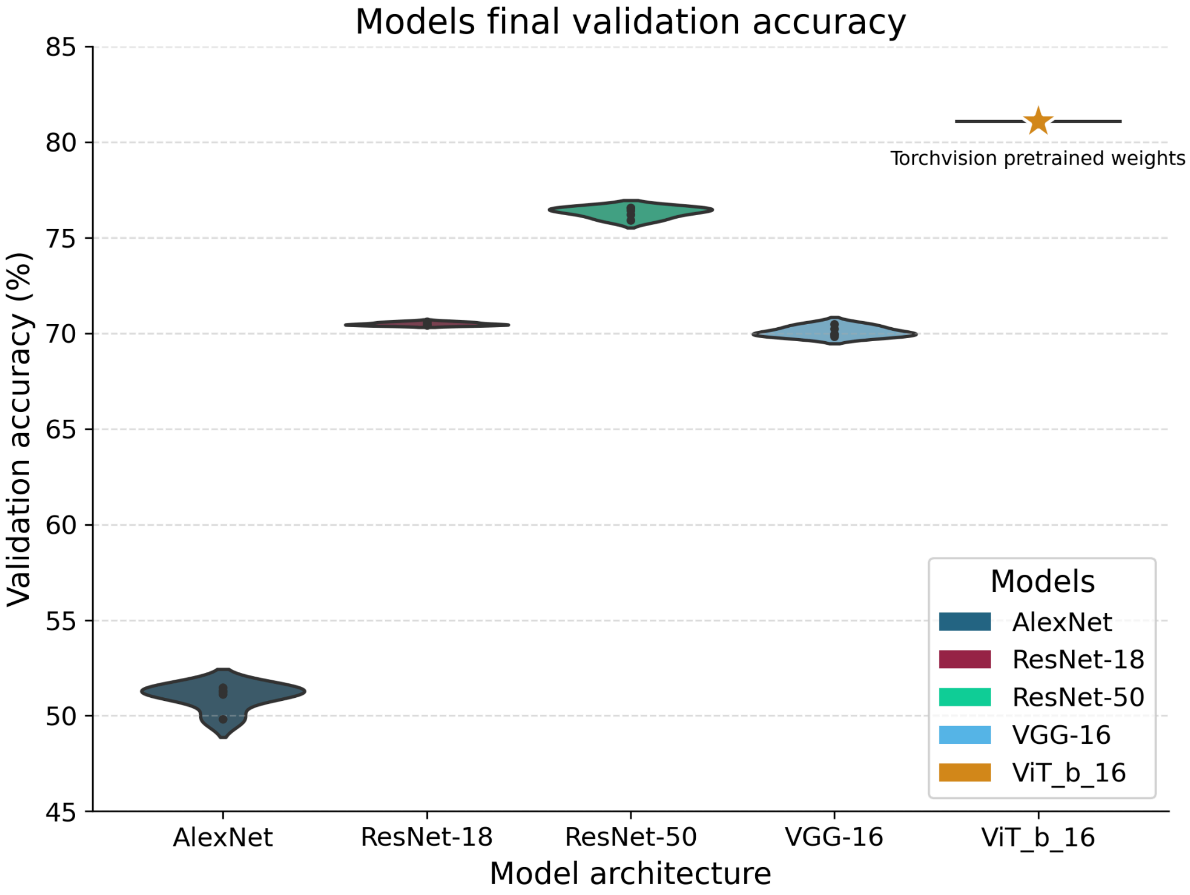


Figure S1. **DNN validation accuracy. Related to Figure 2.** Final ImageNet ILSVRC 2012 validation accuracy for each architecture and random seed. AlexNet achieved the lowest accuracy (~51%), followed by ResNet-18, ResNet-50, and VGG-16 (~70–76%), with ViT-B/16 achieving the highest accuracy (~81%). Each dot represents one model initialization (5 seeds per convolutional architecture; 1 initialization for the pretrained ViT-B/16). Box plots show distributions across seeds. Accuracy values reflect standard object recognition performance and confirm that the trained models span a meaningful range of recognition abilities. Crucially, higher recognition accuracy did not predict higher DNN-EEG alignment, consistent with prior reports [S1,S2].

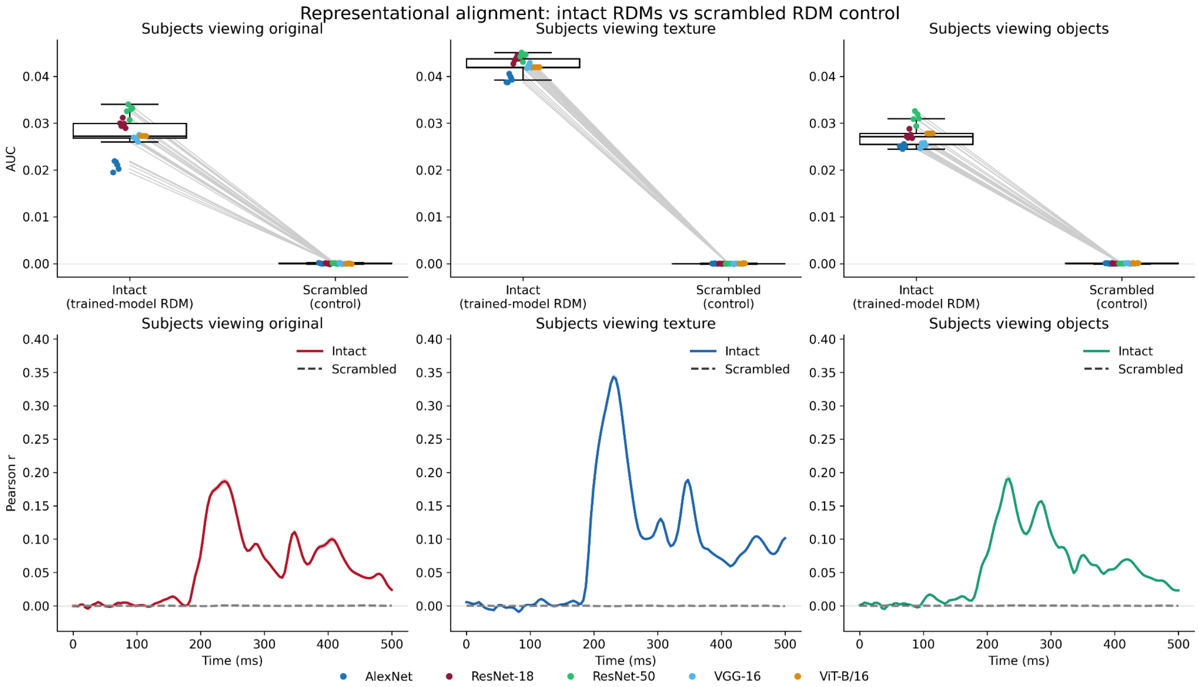


Figure S2. **Control analysis comparing intact versus scrambled RDMs. Related to Figure 2.** We performed a control analysis to test whether the texture advantage is an artifact of overall representational-space volumes by comparing intact trained-model RDMs against randomly scrambled RDMs. The top panels display the Area Under the Curve (AUC) distributions, while the bottom panels track the correlation over time. The plotted dashed-line for the scrambled RDMs is averaged across all permutations. The scrambled RDMs failed to reproduce the original alignment pattern, confirming that the high DNN-EEG alignment for texture-synthesized images is a real effect rather than a difference in the distribution of dissimilarity values.

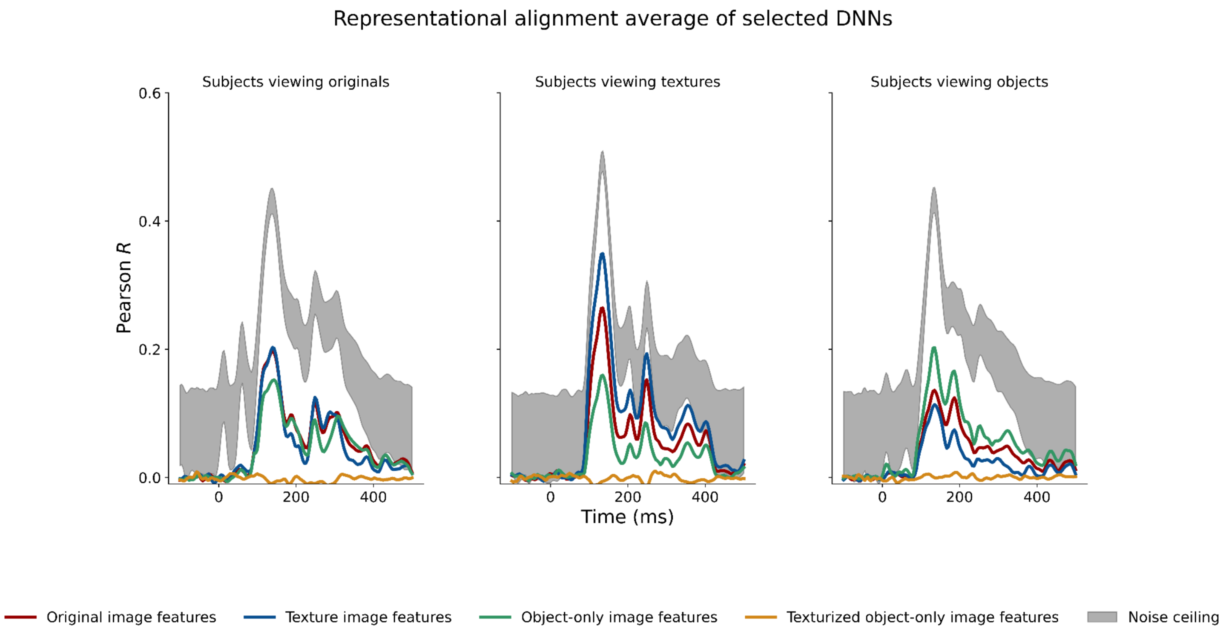


Figure S3. **DNN-EEG alignment using texture-synthesized object-only features. Related to Figure 3.** To test whether DNN-EEG alignment can be explained by the local texture of the isolated object alone, we generated texture-synthesized object-only images, then extracted DNN features from them. These images preserve the local texture-like statistics of the isolated object while removing background and contextual information and disrupting global object form. We used these features to predict EEG responses in all three viewing conditions. Time-resolved alignment (Pearson *r*, averaged across all five architectures) is shown for each viewing condition (subjects viewing originals, texture-synthesized, and object-only images); colored lines indicate the DNN feature set used for prediction (red = original, blue = texture-synthesized, green = object-only, orange = texture-synthesized object-only), and gray shaded regions show the upper and lower noise ceiling bounds. Texture-synthesized object-only features align poorly across all conditions, indicating that DNN-EEG alignment is not explained by isolated object texture but instead depends on broader, full-image local statistics that include background and contextual structure. The above-zero alignment for object-only features, together with the decoding results in Figure S4, is consistent with object-related information being present in the neural signal.

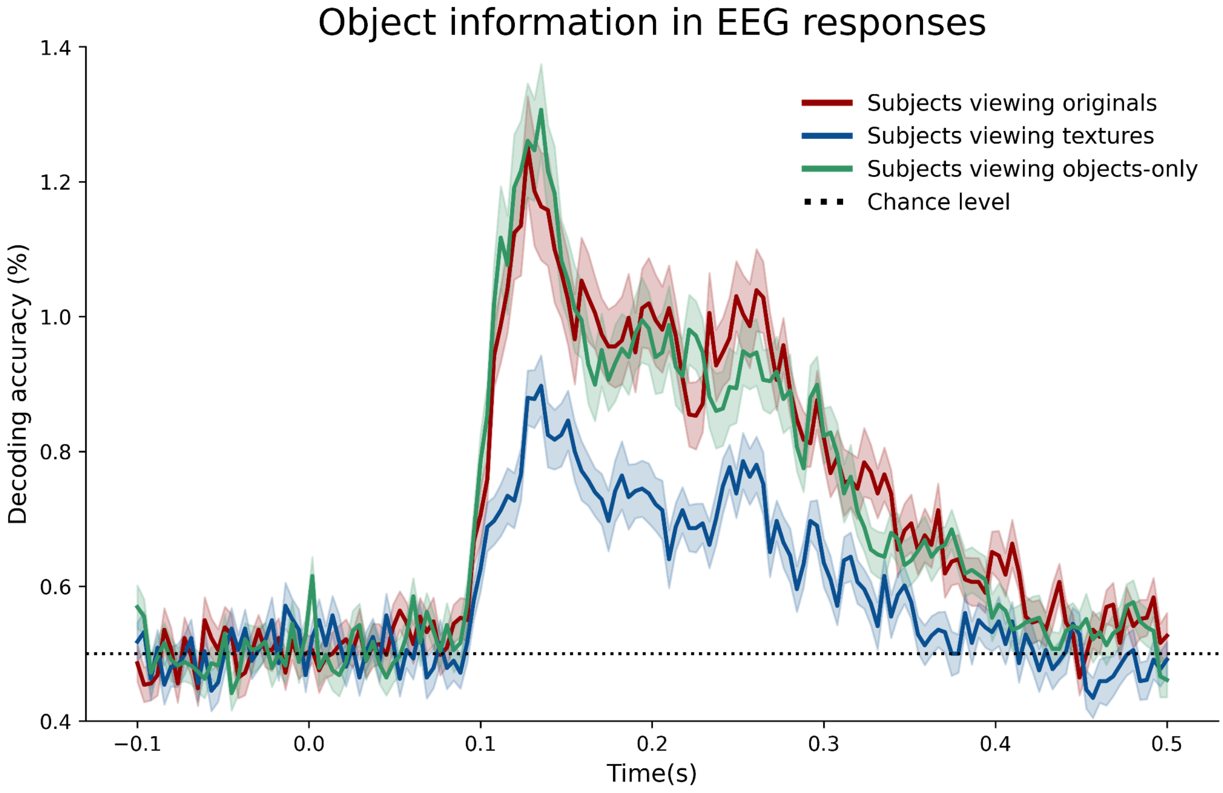


Figure S4. **Object category decoding across image conditions. Related to Figure 4.** Time-resolved SVM decoding accuracy (%) for 200 object categories from single-trial EEG patterns (17 posterior electrodes) for each image condition: original natural scenes (red), texture-synthesized (blue), and object-only (green). Shaded regions: ±1 SEM across participants (n=52). Horizontal dashed line: chance level (0.5%, 1/200 categories). Decoding exceeded chance-level for all three conditions, with natural and object-only conditions reaching higher peak accuracy (1.25% and 1.31%, respectively) than texture-synthesized (0.90%), consistent with greater object form information in the former conditions. Notably, texture-synthesized decoding, while lower, remains reliably above chance, reflecting object category information encoded in local image statistics. The presence of reliable object information across conditions confirms that EEG measurements were sensitive to object-related signals, and that weaker DNN-EEG alignment for natural and object-only conditions is unlikely to reflect a measurement limitation.

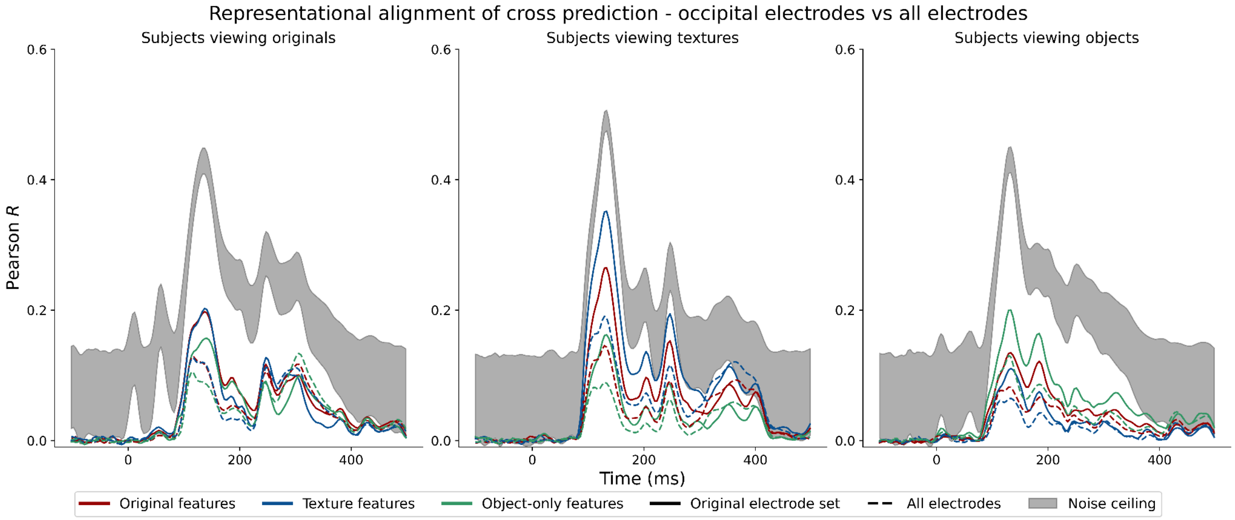


Figure S5. **Representational alignment using posterior versus all electrodes. Related to Figure 3.** DNN-EEG alignment for the three image conditions (original natural scenes, texture-synthesized, and object-only), computed using the 17 posterior electrodes used in the main analyses and, for comparison, using all 64 recorded electrodes. Solid lines show the original 17 posterior electrodes analysis; dashed lines show the all 64 electrodes analysis. Gray shading shows the all electrode noise ceiling bounds. Including all electrodes attenuated the overall alignment but preserved the ranking across conditions, with texture-synthesized images yielding the strongest alignment in both electrode sets. This attenuation is consistent with the relevant signal being concentrated in posterior visual sensors; the all-electrode noise ceiling is correspondingly lower, reflecting the reduced explainable variance introduced by including non-visual sensors. The preserved ranking indicates that restricting the main analyses to posterior electrodes did not produce the texture advantage.
